# Components of Isolated Skeletal Muscle Differentiated Through Antibody Validation

**DOI:** 10.1101/2023.05.20.541600

**Authors:** Dominique C. Stephens, Margaret Mungai, Amber Crabtree, Heather K. Beasley, Edgar Garza-Lopez, Larry Vang, Kit Neikirk, Zer Vue, Neng Vue, Andrea G. Marshall, Kyrin Turner, Jian-qiang Shao, Bishnu Sarker, Sandra Murray, Jennifer A. Gaddy, Jamaine Davis, Steven M. Damo, Antentor O. Hinton

## Abstract

Isolation of skeletal muscles allows for the exploration of many complex diseases. Fibroblasts and myoblast play important roles in skeletal muscle morphology and function. However, skeletal muscles are complex and made up of many cellular populations and validation of these populations is highly important. Therefore, in this article, we discuss a comprehensive method to isolate mice skeletal muscle, create satellite cells for tissue culture, and use immunofluorescence to validate our approach.

**Graphical Abstract:** 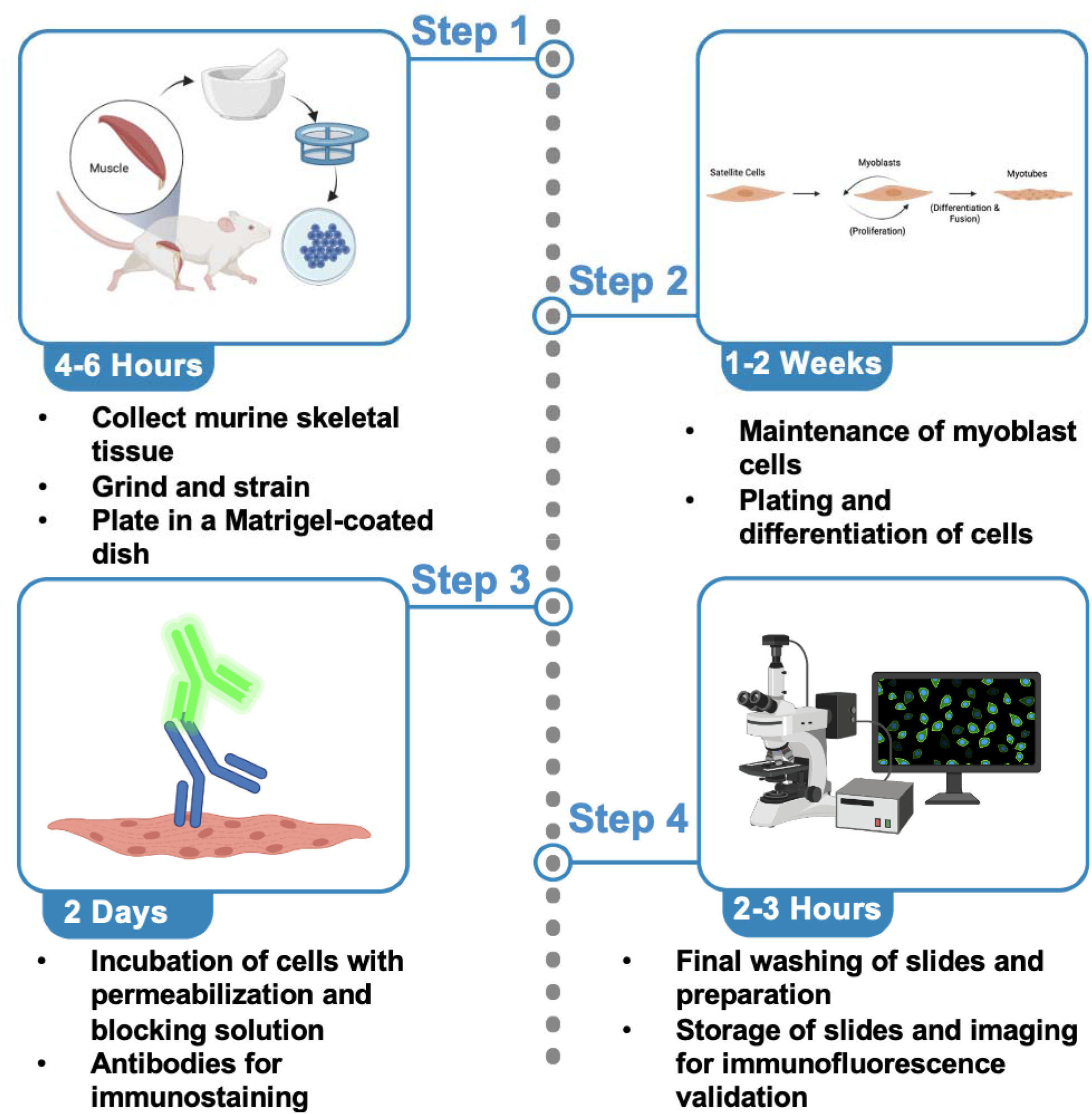

## Before you Begin

Skeletal muscles (SkM) allow for animals and humans to be mobile ^1^, serving many important roles and constituting nearly half of the total mass of the adult human body ^2^. Defects in skeletal muscle mass can cause atrophy and other pathological diseases ^3^. Beyond only mediating glucose uptake in an insulin-dependent manner, skeletal muscle also plays important roles in the metabolism and development of diabetes ^4^. Since the first description of skeletal muscle diseases ^5^, there have been numerous discoveries describing their pathology and the next step in studying these pathologies is characterizing the different cellular populations residing within them. Isolating cells from these muscles allows for models to develop more complex studies to understand how these pathological mechanisms work. In addition to muscle diseases, skeletal muscles are also used to study immunological, neuronal, and other chronic diseases ^6^. While past studies have used immortalized myogenic cells, myoblasts offer unique advantages to understanding the process of myogenesis, which is an avenue for the repair of injured myofibers ^7^. Specifically, skeletal muscle cells are essential for studies on exercise and insulin stimulation. They are also useful experimental models to answer more complex questions, such as the effects of insulin stimulation ^8^ on organelle morphology and the efficacy of new microscopy methods like Focused Ion Beam Scanning Electron Microscopy (FIB-SEM)^9^. Yet, protocols that allow for the differentiation and isolation of myotubes and myoblasts remain limited.

Here we offer two aims, firstly to show how to develop isolate myoblasts, or differentiated myotubes, from murine skeletal muscle (**Figure 1**). Secondly, developing antibody-based approaches for validating SkM cells has been a challenge. Here we also offer a technique for myoblast validation. Antibodies are useful for validating different populations of skeletal muscle cells. Antibodies allow researchers to study the diversity of muscle fibers and cells while providing important insights into cellular processes and disease development. Here, we listed common antibodies used to study different cell populations in SkM tissue (**Table 1**).

**Figure 1:**
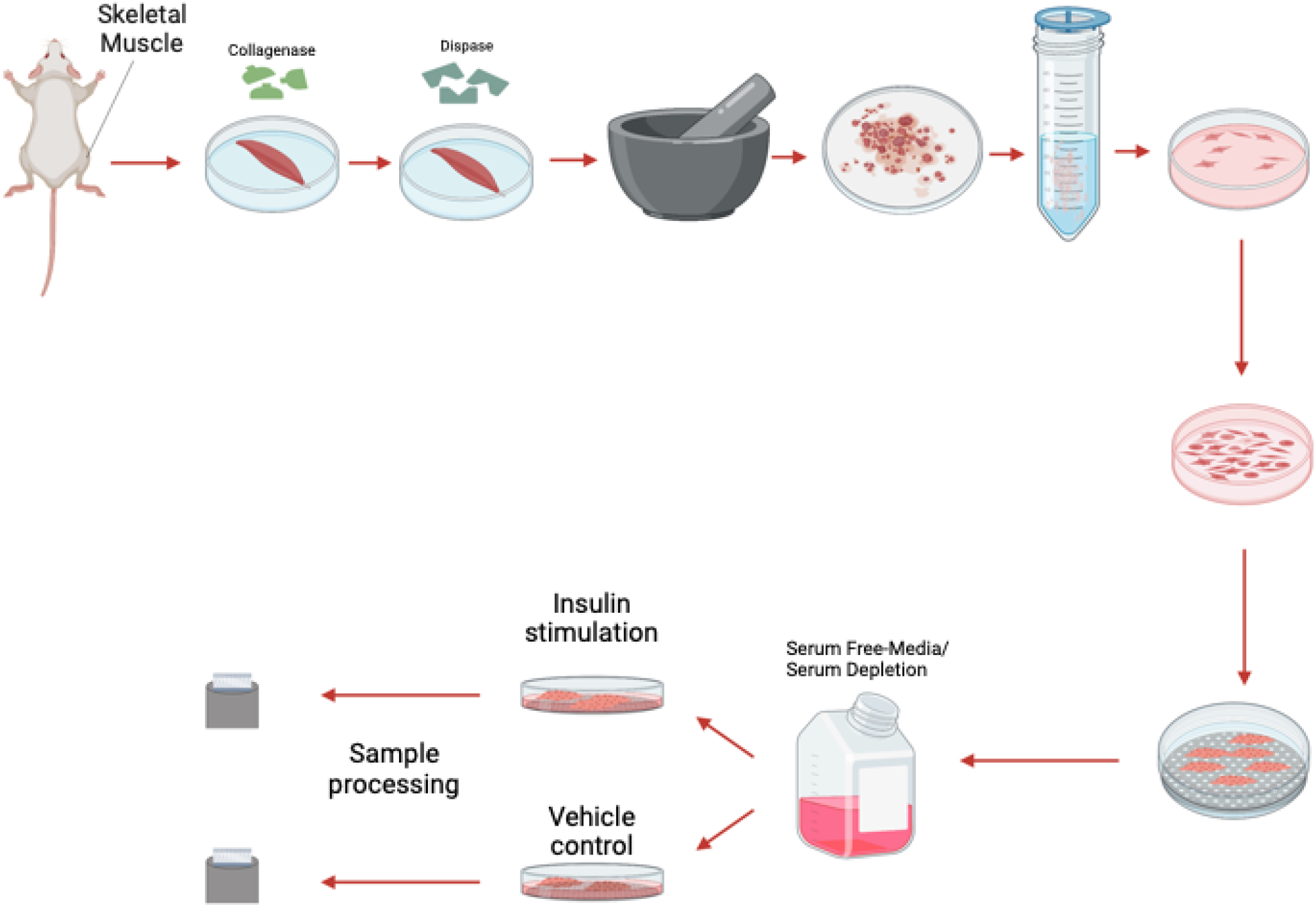
The process of myoblast isolation from gastrocnemius muscle.

**Table 1:**
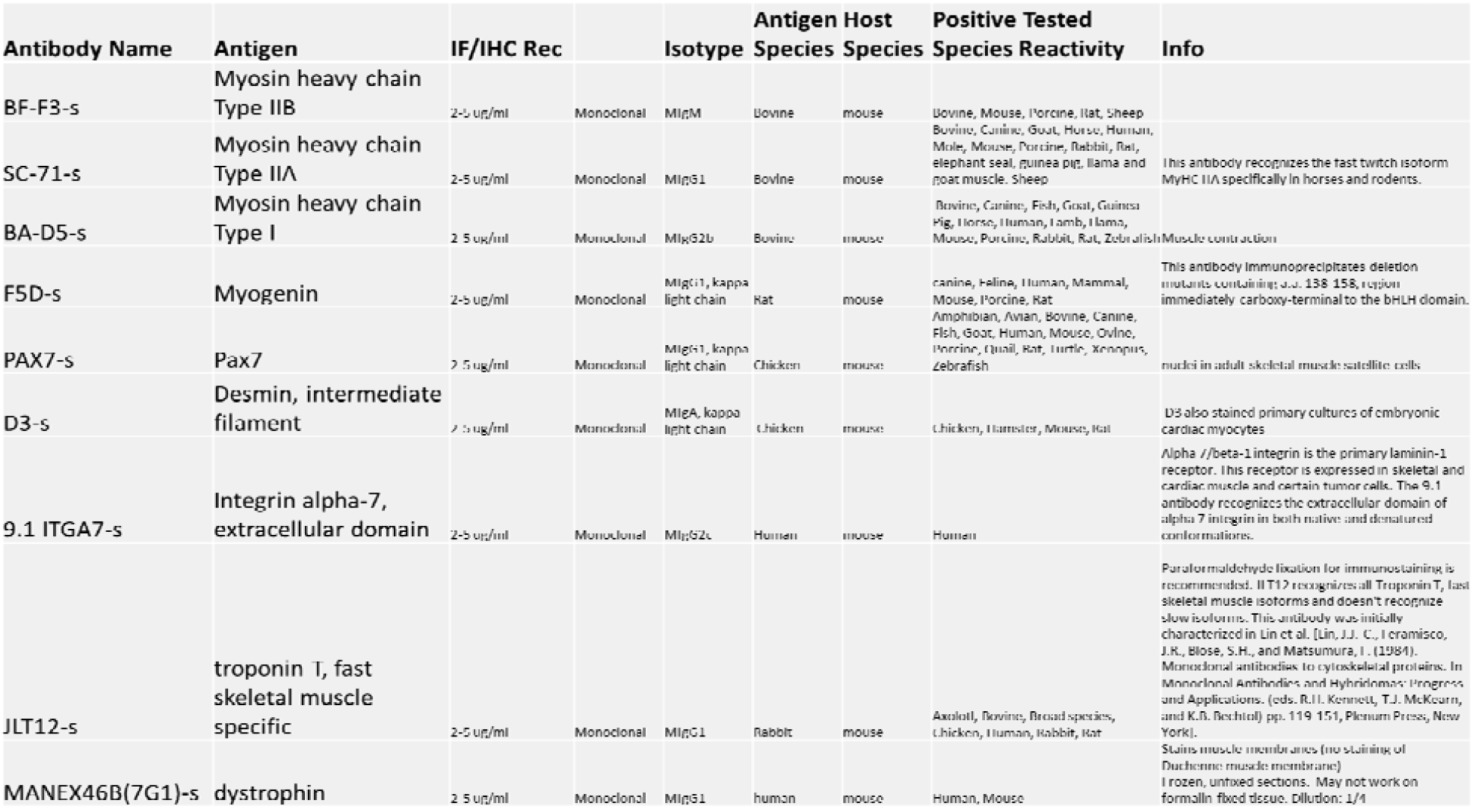
A list of antibodies and their respective antigens for skeletal muscle validation after isolation. The Table was obtained from https://dshb.biology.uiowa.edu/.

SkM tissue is composed of various cell types with different functions, including myoblasts and fibroblasts ^3^. Skeletal myoblasts drive muscle regeneration after injury, while fibroblasts create extracellular matrix components and secrete growth factors ^10^ (**Figure 2**). Morphologically, fibroblasts are larger than myoblasts and contain more vesicles ^11^. Beyond this, while mononuclear cells replicate, as they form sheets of multinucleated myotubes, proliferation is impaired and myogenin is elevated ^12^. Myoblasts’ process of differentiation mimics that of *in vivo* myogenesis, with the structure of myoblasts affecting that of differentiated myotubes ^13^. Given that these populations have morphological differences ^14^, validating the myoblast or myotube stage is of critical importance, especially for experiments that seek to study homogenous populations and fine ultrastructural changes. Here, we also present how antibodies and fluorescence light microscopy can be used to validate different cell populations in skeletal muscle tissue. However, before you begin care should be taken in experimental design and selection for which populations you want to obtain. Here, we propose a standardized approach to isolate and identify different skeletal muscle cell populations. Using these methods, we looked at the effects on insulin stimulation on oxygen consumption rate (OCR) and have verified past studies which have implicated changes in respiration following insulin stimulation in a potential optic atrophy protein-1 (OPA-1) mediated manner ^15^.

**Figure 2:**
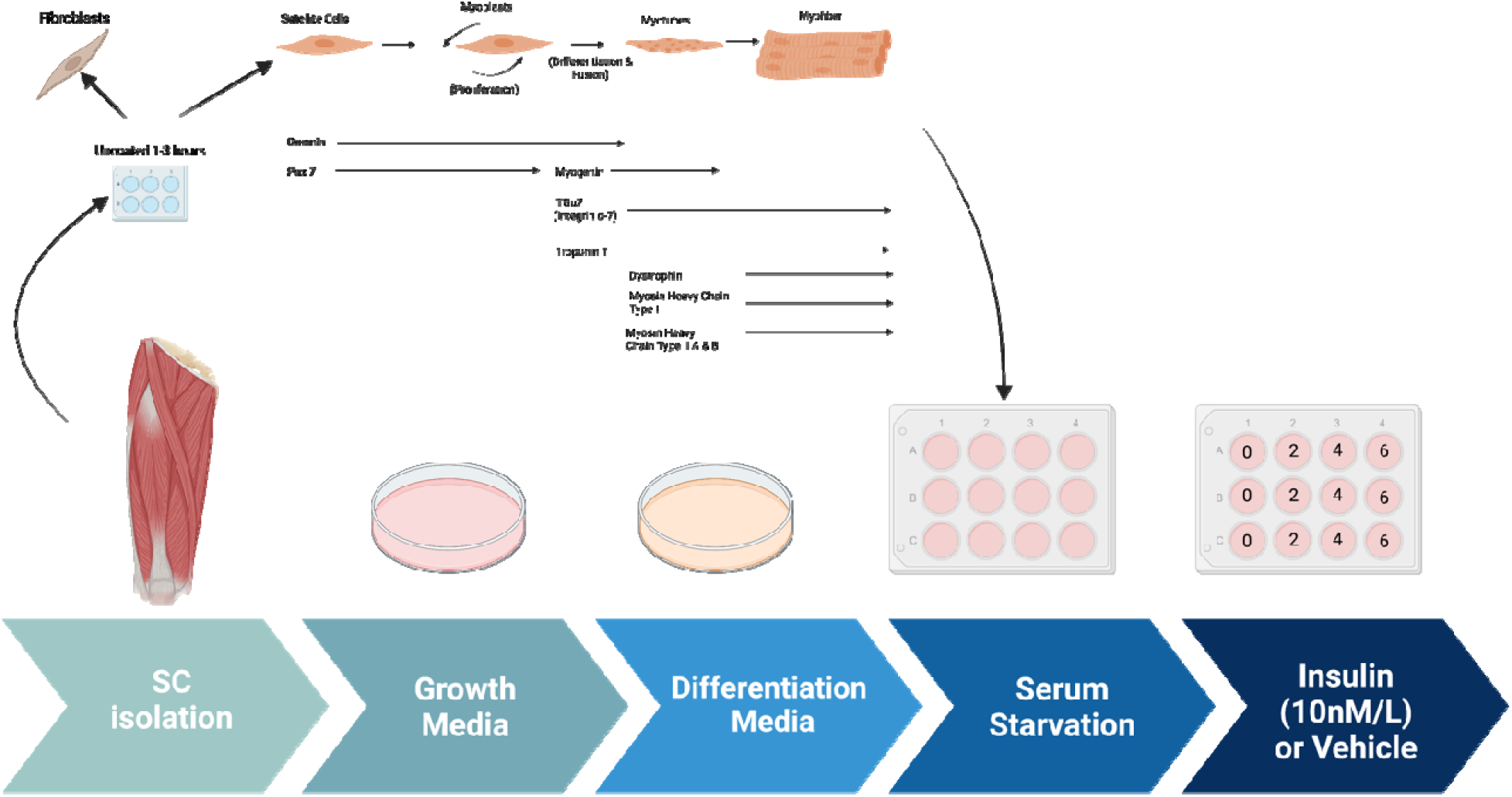
The process of myotube differentiation from myoblasts and utilization for serum.

## Methods and Materials

### Animal Studies

All mice utilized had a C57Bl/6J background. Mice studies followed previous studies with ^16–18^ with weaning at 3 weeks of age and maintained on standard chow (2920X Harlan Teklad, Indianapolis, IN, USA), and at 22°C with a 12 hr light, 12 hr dark cycle with free access to water and standard chow. All mouse experiments were conducted in alignment with the animal research guidelines from NIH and were approved by the University of Iowa IACUC.

### Light Microscopy

Staining of myotubes was performed on Olympus IX-81. Myotubes were plated on 35 mm dishes with a glass bottom and imaged for light microscopy. Transmission electron microscopy performed with Joel 1400, operating at 80 kV, as we have done previously ^17^. 3 days after differentiation, myotubes were infected with GFP-expressing adenovirus per previous procedures ^16^. All imaging experiments were performed at Central Microscopy Research Facility, University of Iowa, while adenoviruses were obtained from University of Iowa Viral Vector Core facility.

### Seahorse Analyzer

Mice were anesthetized using a mixture of 5% isoflurane/oxygen. Oxygen consumption rate was measured for using an XF24 bioanalyzer (Seahorse Bioscience: North Billerica, MA, USA), as previously described ^16,19^. Myotubes and myoblasts were plated at a density of 20 × 10^3^ per well and differentiated for 3 days. Isolated myotubes and myoblasts were either treated with 10 nmol/L insulin for the specified time ^15^. Media was replaced with XF□DMEM (supplemented with 1 g/L D-Glucose, 0.11 g/L sodium pyruvate, and 4 mM L-Glutamine) and cells were deprived of CO_2_ for 60 minutes. Oligomycin (1 μg/ml), carbonyl cyanide 4- (trifluoromethoxy)phenylhydrazone (FCCP; 1 μM), rotenone (1 μM), and antimycin A (10 μM) treatment occurred.

For quantifications, time points 1-3 measure basal respiration, or baseline rate of oxygen consumption by cells in culture without any treatment. Oligomycin (1 μg/ml) was added to inhibit ATP synthase, which reduces mitochondrial respiration and leads to an increase in proton gradient, to measure the amount of oxygen consumed by the myoblasts and myotubes to maintain the proton gradient in time points 4-6. carbonyl cyanide 4- (trifluoromethoxy)phenylhydrazone (FCCP; 1 μM) was then added in time points 7-9 which allows electrons to flow freely through the electron transport chain allowing for measurements of reserve capacity and maximum oxygen consumption. Finally, rotenone (1 μM) and antimycin A (10 μM) were added in time points 10-12 which inhibit electron transfer from NADH to ubiquinone and ubiquinol to cytochrome c, respectively, to measure non-mitochondrial respiration ^20^.

For normalization of proteins, after measurement 20 μl of 10 mM Tris with 0.1% Triton X-100 was added at pH 7.4 to lyse cells per prior protocols ^19^, and media was replaced with 480 μl of Bradford reagent.

### Transmission Electron Microscopy

Myoblasts and myotubes were isolated according to the step-by-step below and placed in six-well poly-D-lysine-coated plates for TEM processing per established protocols ^1,2^. Briefly, cells were fixed by incubating at 37 °C with 2.5% glutaraldehyde in 0.1 m sodium cacodylate buffer for one hour, then rinsed twice with 0.1 m sodium cacodylate buffer, prior to fixation at room temperature for 30 min to 1 h using 1% osmium tetroxide and 1.5% potassium ferrocyanide in 0.1 m sodium cacodylate buffer. Samples were washed for 5 min with 0.1 m sodium cacodylate buffer (7.3 pH), then diH2O (2 × 5 minutes). Samples were incubated with 2.5% uranyl acetate, diluted with H2O, at 4 °C overnight. Samples were dehydrated and ethanol was replaced with Eponate 12 mixed in 100% ethanol in a 1:1 solution for 30 min at RT. This was repeated three times for 1 h using 100% Eponate 12, the media was replaced and plates were cured in an oven at 70 °C overnight.

After cracking and submerging the plate in liquid nitrogen, an 80 nm thickness jeweler’s saw was used to cut the block to fit in a Leica UC6 ultramicrotome sample holder. From there, the section was placed on formvar-coated copper grids. These grids were counterstained in 2% uranyl acetate for 2 min and Reynold’s lead citrate for 2 min. Images were acquired by TEM on either a JEOL JEM-1230, operating at 120 kV, or a JEOL 1400, operating at 80 kV. Analysis performed via protocols by Lam et al. (2021) ^1^.

### Key Resource Table

**Key Resource Table:**
| REAGENT or RESOURCE | SOURCE | IDENTIFIER |
| --- | --- | --- |
| 4% paraformaldehyde | ThermoFisher | J61899-AK |
| Beta-Mercaptoethanol | Gibco | 21985-023 |
| Collagenase II | Gibco | 17101015 |
| DMEM (+ 4.5 g/L D-glucose, + L-Glut, - Sodium Pyruvate) | Gibco | 11965-092 |
| F-12 (+L-Glut) | Gibco | 11765-054 |
| FBS | Atlanta Biologicals | S11550 |
| Fungizone | Gibco | 15290-018 |
| Human FGF-basic (FGF-2/bFGF) Recombinant Protein | ThermoFisher | 13256-029 |
| Insulin-transferrin-selenium-X (100x) | Gibco | 41400045 |
| MEM Non-essential Amino Acids | Gibco | 11140 |
| PBS, pH 7.4 | Gibco | 10010023 |
| Penicillin-Streptomycin | Gibco | 15140 |
| StemPro™ Accutase™ | ThermoFisher | A1110501 |
| Triton X-100 (1%) | Gibco | HFH10 |
| **Quantification and Software** |  |  |
| GraphPad | GraphPad Software, San Diego, California USA | www.graphpad.com |
| Image J | Schindelin et al.5 | https://imagej.net/ |

Prior to the protocol, make the following reagents and solutions and sterilize all equipment in an autoclave.

### 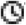 Timing: 1 hr

Initial PBS Wash mixture:

| Name | Volume |
| --- | --- |
| PBS | 25 mL |
| Fungizone | 75 µL |
| Penicillin-Streptomycin | 250 µL |

Initial DMEM-F12 incubation mixture:

| Name | Volume |
| --- | --- |
| DMEM-F12 | 250 mL DMEM (+ 4.5 g/L D-glucose, + L-Glut, - Sodium Pyruvate) + 250 mL F-12 (+L-Glut) |
| Collagenase II | 1300 mg |
| Penicillin-Streptomycin | 6.4 mL |
| Fungizone | 2.0 mL |

Secondary DMEM-F12 incubation mixture:

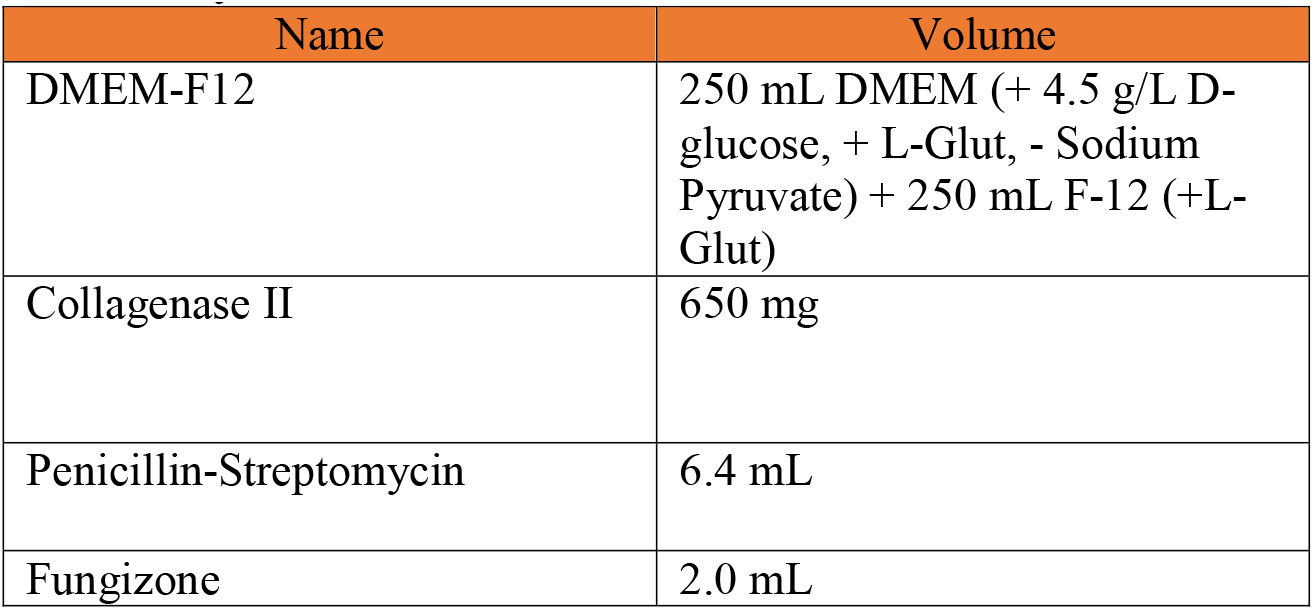
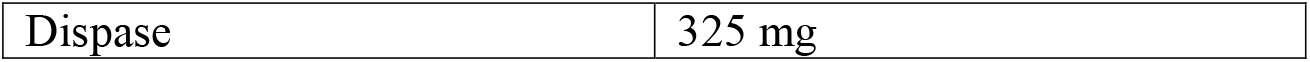

DMEM-F12 Growth Media:

Mix the following. Use a sterile filter with Millipore brand 0.22 uM filter units. Store at 4C for no longer than 2 months. Add bFGF (10ng/mL) to the aliquot just before adding it to plate.

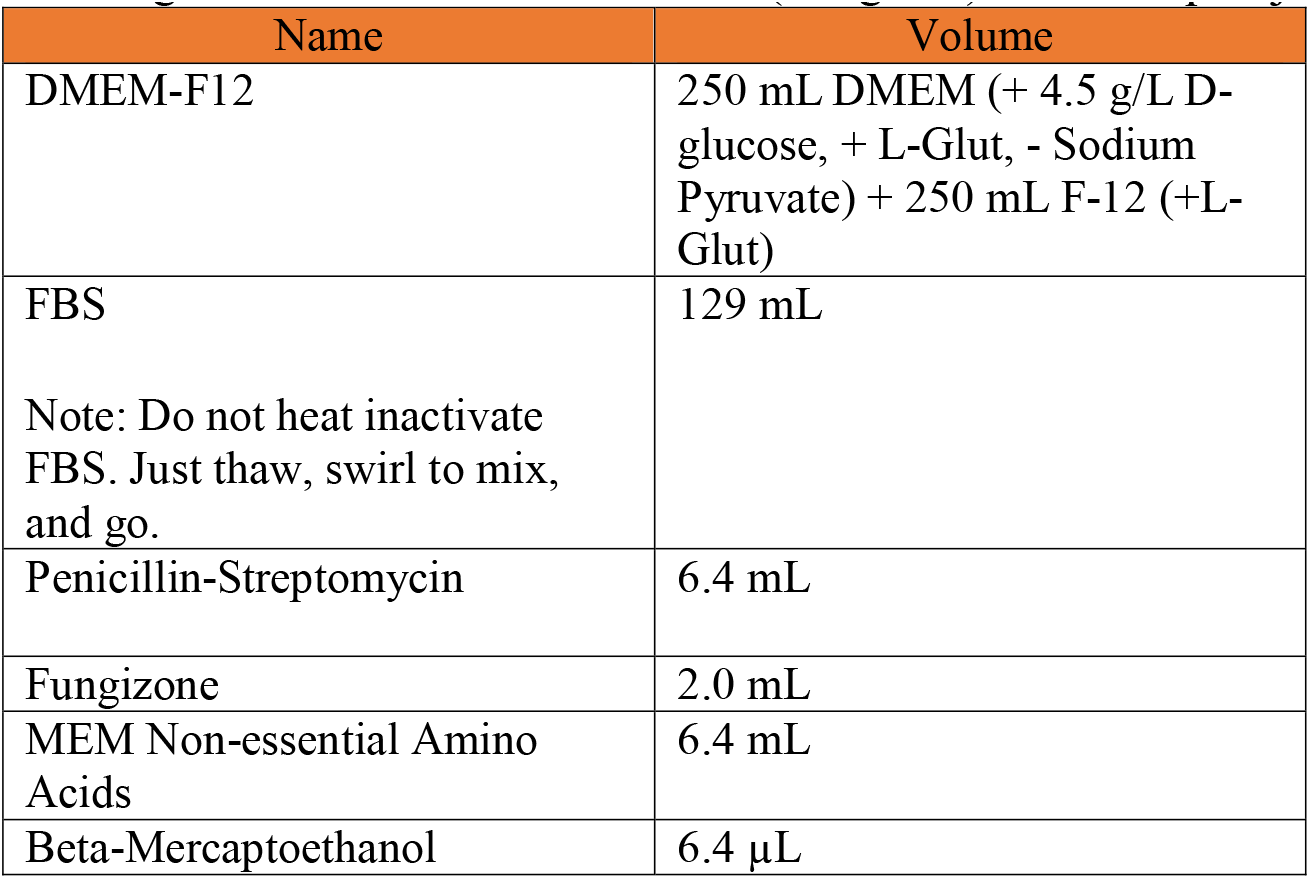

Permeabilization Buffer:

| Name | Volume |
| --- | --- |
| PBS | 495.5 mL |
| Triton X-100 | 0.5 mL |

Differentiation medium:

| Name | Volume |
| --- | --- |
| DMEM (+ 4.5 g/L D-glucose, + L-Glut, - Sodium Pyruvate) | 250 mL |
| F-12 (+L-Glut) | 250 mL |
| FBS | 10.5 mL |
| Note: Do not heat inactivate FBS. Just thaw, swirl to mix, and go. |  |
| Insulin-transferrin-selenium-X (100x) | 5.3 mL |

Reconstitute Human FGF-basic (FGF-2/bFGF) Recombinant Protein (here we use ThermoFisher 13256-029). Briefly, to prepare a stock solution of bFGF at a concentration of 0.1 mg/mL, reconstitute it in 100 μL of 10 mM Tris (pH 7.6). Dilute in buffer containing 0.1% BSA and store in polypropylene vials for up to six months at -20°C. Make aliquots to avoid repeated freezing and rethawing.

### Step-by-Step

#### Myoblast Isolation: Removal of Muscle

The below offers a basic text for the isolation of myoblast.

***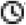 Timing: 30 minutes***.

1. Following IACUC guidelines, euthanize mice.
2. Collect muscle tissue from the gastrocnemius, quadriceps, soleus, and hamstring muscles in both legs at 4-8 weeks of age from 4-6 mice.

**NOTE:** Less or more mice can be utilized but we generally found batches of 4-6 to be a good quantity.

**CRITICAL:** This protocol works to make tissue-type specific cell lines. If the muscle types will not be combined, double the number of mice, to 8-12 mice, to ensure an adequate sample count.

3. Transfer the collected tissue to an Eppendorf tube.

**NOTE:** From this point on, ensure a sterile environment is utilized.

#### Myoblast Isolation: Preparation of muscle, shaking, and grinding of tissue

***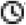 Timing: 2.5 hrs***.

4. Wash isolated tissue 2-3 times with the initial PBS wash mixture.

**NOTE:** The PBS solution is prepared right before dissecting the tissue.

5. Incubate muscle tissue in the initial DMEM-F12 incubation mixture.

**CRITICAL:** Avoid filtering the DMEM-F12 media containing collagenase, 1% pen/strep, and 3 μL/mL Fungizone. This initial DMEM-F12 incubation mixture must be chilled (4 °C) when added to reduce temperature shock.

6. Maintain the muscle solution in a 37 °C water bath for 10-15 mins.
7. Shake at 220 rpm, for an overall time of 1.5 hrs.
8. After incubation, wash the tissue 3-4 times with PBS.
9. Incubate in warmed secondary DMEM-F12 incubation mixture while the tissue is shaken for 30 mins in a 37 °C water bath.

**NOTE:** Secondary DMEM-F12 incubation mixture has to be pre-warmed to 37°C to ensure efficient mixing of dispase and since the muscles were at 37°C after incubation.

10. After shaking, grind tissue with a mortar and pestle in the presence of liquid nitrogen.
11. Pass through a 100 μm, then 70 μm, cell strainer.
12. Centrifuge the solution at 1000 rpm for five mins to pellet the cells.

#### Myoblast Isolation: Plating

***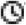 Timing: 1-3 hrs***.

13. Transfer the to a plate and resuspended using DMEM-F12 growth media supplemented with 40 ng/mL bFGF.
14. Pre-plate the cells for 1-3 hours on UNCOATED dishes to reduce the number of fibroblasts.

**CRITICAL:** Fibroblasts can dilute satellite cells. Recommended for dystrophic or injured muscle. Pre-plating on an uncoated plate causes fibroblasts to stick and be isolated. Fibroblasts can separately be used to isolate and for other experiments.

15. Dilute cells 1:15 in PBS, then plate in a Matrigel-coated dish.

**NOTE:** To create Matrigel-coated dishes, dilute stock concentration (while keeping on ice) to 1:15 in sterile PBS in the hood. Put Matrigel solution on flask/plate, shake/tilt to coat the bottom, incubate at room temperature in a chemical hood for 30 mins, and remove Matrigel solution back into its original tube. Matrigel solution may be reused up to 5 times in total.

#### Myoblast Isolation: Differentiation

The below protocol offers subsequent differentiation to myotubes, if desired.

***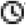 Timing: 1-2 weeks***.

1. Wait for activation, which takes 24-48 hrs, after which myoblasts will grow rapidly.
  a. To maintain healthy myoblast cells, use the growth media supplemented with bFGF (10 ng/mL).

**CRITICAL:** Use Differentiation Medium to go from myoblasts to myocytes and then to myotubes (Figure 2).

2. Plate primary myoblast at ∼.8 × 10^6^ cells per well and to differentiate the cells, add differentiation media, supplemented with 1:10,000 bFGF.

**NOTE:** This will depend on # of cell passages and type of treatment, adjust accordingly.

3. Incubate for 4 to 7 days for differentiation to myotubes.

**NOTE:** Switch out with fresh differentiation media every 2 days, supplemented with 1:10,000 bFGF.

4. Cells are split using 2-5 mL of accutase for 5-15 minutes, dependent on cell count.

**Note: DO NOT** use trypsin to split the cells. Accutase is less harsh to the extracellular matrix, surface proteins, and cytoskeleton of skeletal cells than trypsin, so it is highly preferred ^21^. Accutate total incubation time and volume will differ depending on cell yield.

5. Cells are maintained in (5% CO_2_) at 37°C. If growing myotubes, A confluency of 70-85% has to be reached prior to adding growth media.

#### Validation Day 1: Permeabilization and Blocking

Immunofluorescence staining is effective for examining differences in skeletal muscles simultaneously. Refer to Table 1 for a list of validated primary antibodies for skeletal muscles. Select secondary antibodies that are compatible with the epifluorescence or confocal microscope available to you.

***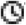 Timing: 1.5 hours***.

**CRITICAL:** All steps are performed at room temperature unless otherwise indicated.

**NOTE:** This protocol for Immunofluorescence staining and antibody validation of isolated skeletal muscle cells is an adaptation of Esper et al., skeletal muscle tissue immunofluorescence labeling protocol ^22^.

1. Fix cells by incubating them in 4% paraformaldehyde (PFA) for five mins.
2. Wash three times for five mins using phosphate-buffered saline (PBS).

**NOTE:** Ice-cold 100% methanol or acetone is an effective fixative for cryosections and more suited for some antigens. Acetone is less harsh than methanol.

3. Incubate cells in permeabilization buffer for 10 mins.
4. Incubate cells in blocking solution for 1 hr at room temperature or overnight at 4 °C.

**NOTE:** When using permeabilization buffer, keep the solution away from the hydrophobic barrier to avoid loss of hydrophobicity. If this happens, wash the slide well with PBS. Include Mouse on Mouse (MOM) blocking reagent at a 1:40 dilution when staining mouse tissue with antibodies raised in the mouse.

#### Validation Day 2: Antibodies

***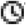 Timing: 30 minutes***.

5. To begin immunostaining, dilute the primary and secondary antibodies in a blocking solution according to the manufacturer’s suggested ratio.

**NOTE**: It is acceptable to dilute antibodies in hybridoma supernatant when targeting multiple antigens.

6. Aspirate the blocking buffer and cover the slide with the primary antibody solution.
7. Incubate the slides overnight at 4 °C.

#### Validation Day 3: Preparing Slides for Imaging

***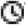 Timing: 2-3 hours***.

8. On the following day, wash three times for five mins with PBS.
9. After washing, cover cells with secondary antibodies diluted in blocking buffer for 1 hr at room temperature in the dark.

**NOTE**: Keep slides in the dark for the remainder of the protocol.

10. After incubation, wash the slides three times for five mins with PBS.
11. Incubate the cells with 1 μg/mL DAPI diluted in PBS for five mins.
12. Wash once with PBS for five min.
13. Aspirate the PBS and place 1–2 drops of mounting media onto the cells
14. Carefully place a coverslip on the slide, while avoiding air bubbles.
15. Let the slides dry in the dark for 1–2 hr before sealing the slides with clear nail polish.
16. Store the slides at 4 °C and image within 2 weeks.

### Expected Outcomes

Upon isolation of myoblast and differentiation into myotubes, we validated their structure in light microscopy (**Figure 3A**). Furthermore, we viewed multinucleated myotubes through TEM to validate the ultrastructure (**Figure 3B**). Transfection further showed myotubes demonstrated fluorescence as expected (**Figure 3C**). From there, we performed staining for myosin and desmin, muscle-specific proteins that play crucial roles in muscle cell structure and function ^23^, to confirm that filaments were present (**Figure 3 D-D’’**).

**Figure 3:**
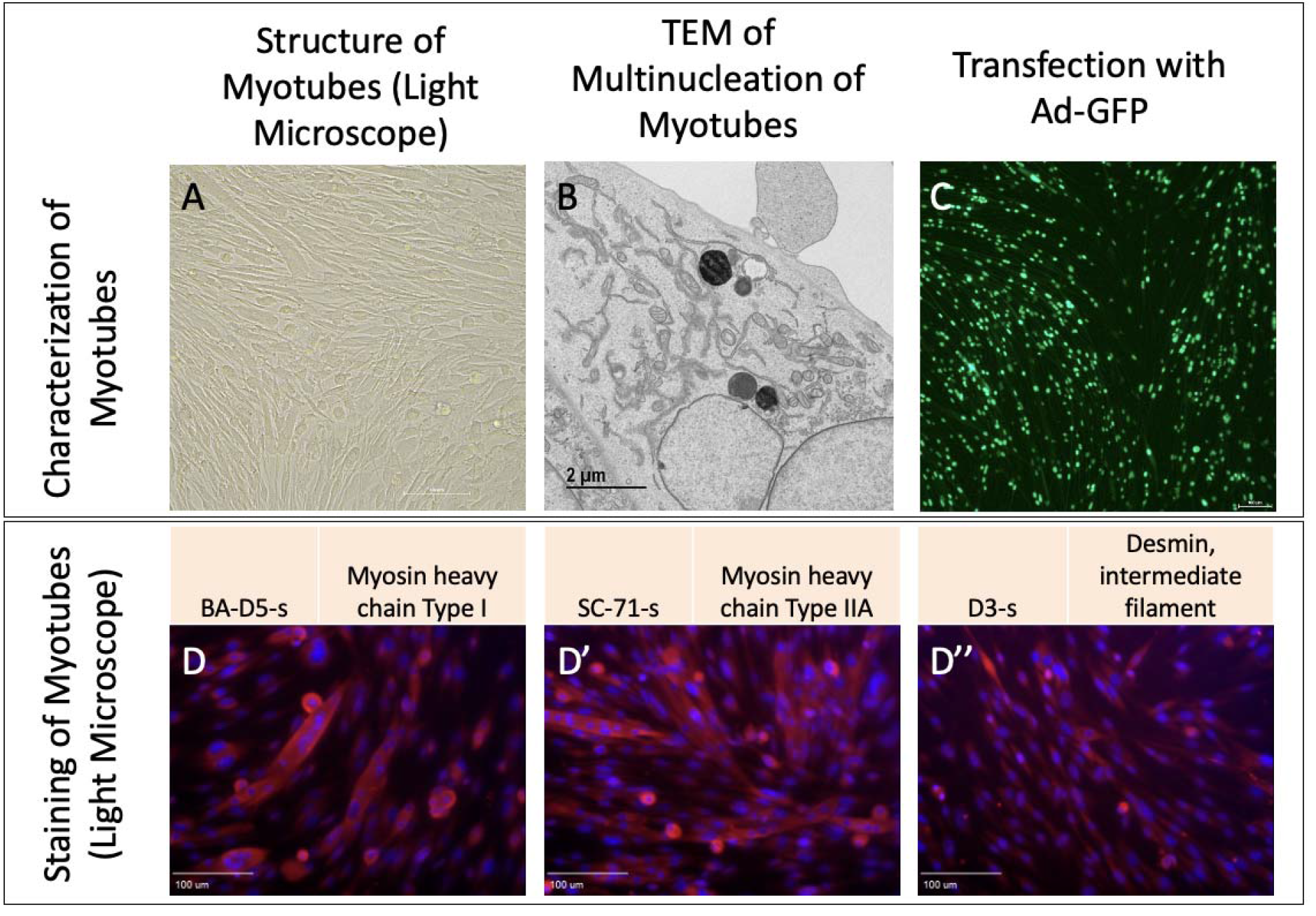
Myotubes as characterized by A. Light microscopy, B. Transmission electron microscopy, and C. Transfection with adenovirus containing the green fluorescent protein gene (Ad-GFP). D. Straining with BA-D5-s, D’ SC-71-s, and D’’ D3-s to show myosin and desmin.

Once myoblasts and myotubes are validated, they can be used for a variety of studies including to measure mitochondrial efficiency with oxygen consumption rate, western blot analysis to look for expression of specific proteins in knockout studies, or a variety of electron microscopy techniques such as serial block-face scanning electron microscopy to perform 3D reconstruction of organelles (**Figure 4**). In the past, following this isolation and differentiation protocol, we have successfully used the protocol by Garza-Lopez et al. (2022) ^24^ for 3D reconstruction and quantification of mitochondria and endoplasmic reticulum, alongside the protocol by Neikirk et al. (2023) ^17^ for 3D reconstruction and quantification of lipid droplets, lysosomes, and autophagosomes. However, from our experience, any validated scanning electron microscopy protocol should be effective following isolation and differentiation.

**Figure 4:**
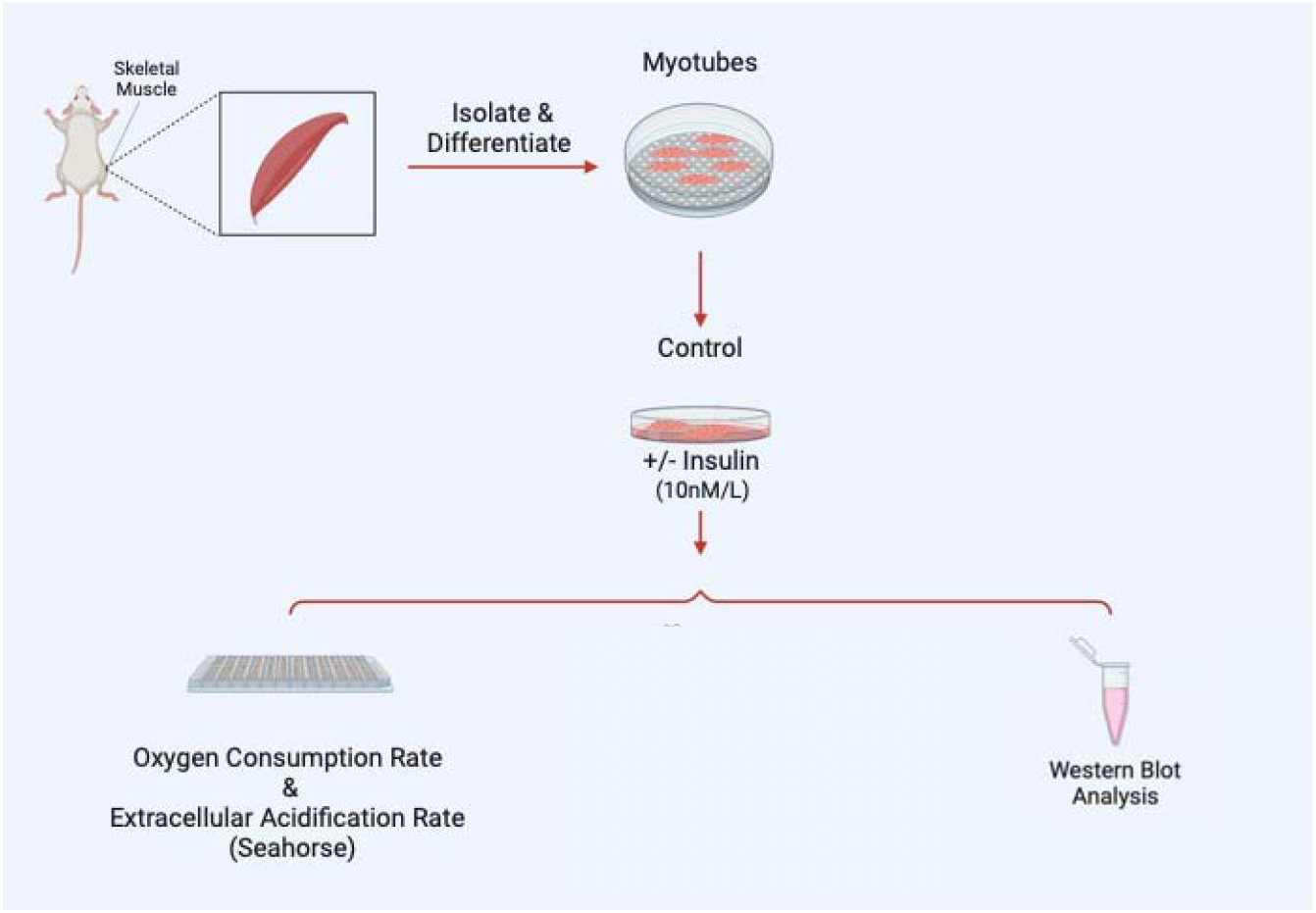
Examples of experiments that may be performed following myotube differentiation and isolation.

As an example, to validate this method, we sought to understand how insulin treatment (10 nM/L) in 2-hour increments may alter myoblast and myotube function through the usage of a Seahorse XF96 analyzer, per past protocols ^25^. We found that for myoblasts, there is a significantly increased basal, maximum, and non-mitochondrial OCR after 2 hours of insulin treatment, while this difference is retained or exacerbated after 4 hours of insulin treatment (**Figure 5A-B**). After 6 hours of insulin treatment, OCR conversely showed significant decreases in all of these parameters (**Figure 5C**). In myotubes, after 2 and 4 hours of insulin treatment, we similarly noted a significant increase in mitochondrial OCR (**Figure 5D-E**). Notably, the increase in basal, ATP-linked, maximum, and non-mitochondrial OCR is much higher in 4 hours than 2 hours. Unlike myoblasts, 6 hours of insulin treatment myotubes did not differ significantly from untreated cells (**Figure 5F**). Importantly, there may be a differential response to insulin treatment in myoblasts and myotubes, highlighting the importance of studying both models. This validated that the function of myoblasts and myotubes are intact following this isolation.

**Figure 5:**
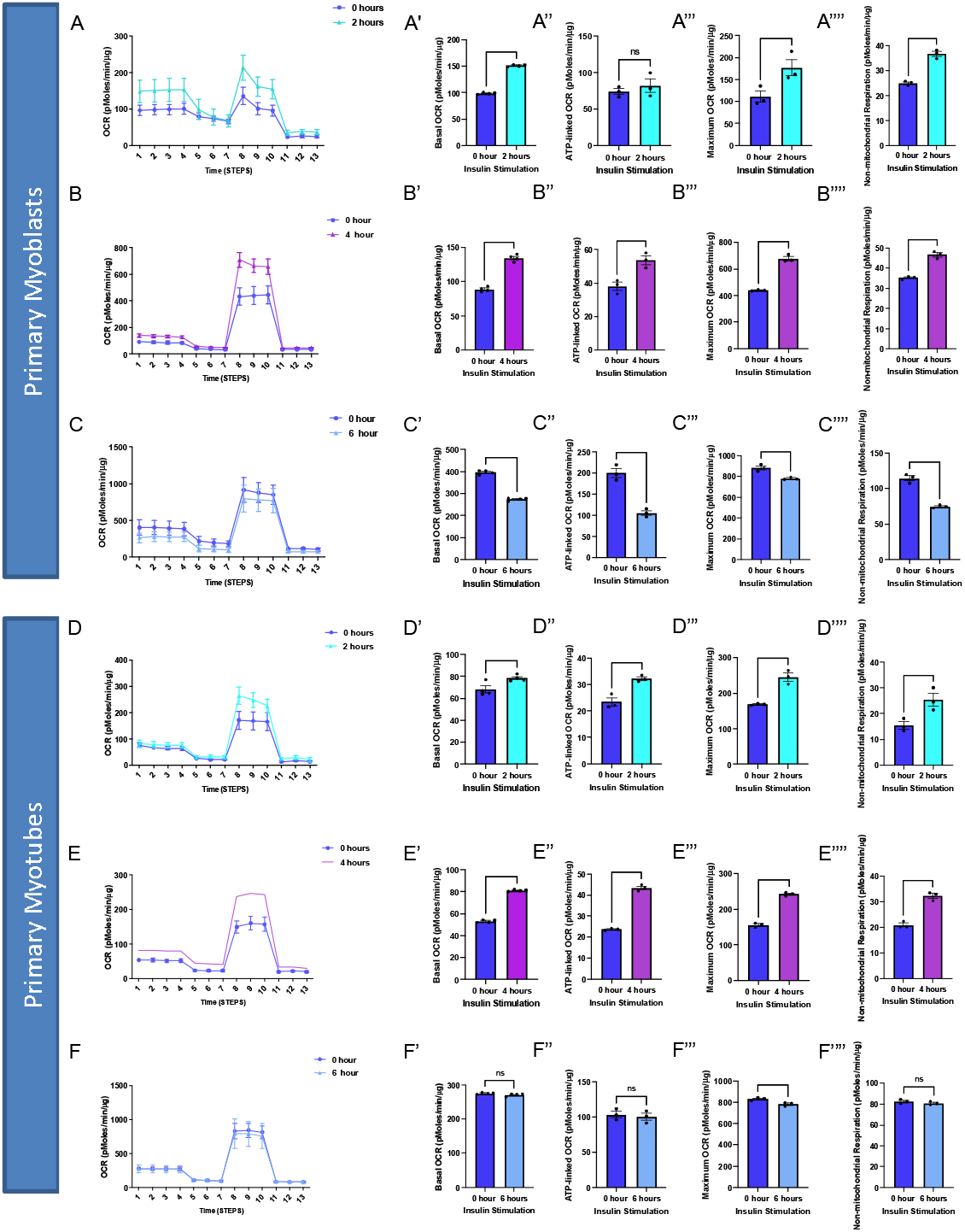
Oxygen consumption rate (OCR) altered in myoblasts and myotubes upon altered insulin stimulation which shows changes in mitochondrial efficiency. (A) Seahorse plot for primary myoblasts following 2 hours of insulin stimulation (B) 4 hours of insulin stimulation and (C) 6 hours of insulin stimulation. (D) Oxygen consumption rate was measured after several inhibitors to measure respiration in primary myotubes after 2 hours, (E) 4 hours, and (F) 6 hours of insulin stimulation. (A’-F’) Basal OCR, which represents respiration under normal, unstressed conditions. (A’’-F’’) ATP-linked OCR, which is respiration associated with ATP synthesis during oxidative phosphorylation, which is marked by a reduction in OCR due to oligomycin. (A’’’-F’’’) Maximum OCR, which is the maximal capacity at which mitochondria may utilize oxygen. (A’’’’-F’’’’) Non-mitochondrial respiration, which can be attributed to factors such as glycolysis or ROS and not due to mitochondrial respiration. These values were compared to the control (blue) in all of these examples. N = 6 per treatment, and * indicates p-value < .05.

From there we sought to elucidate if organelle proteins are affected following insulin treatment and we targeted Optic atrophy protein 1 (OPA-1), which is a mitochondrial inner membrane (IMM) fusion protein that mediates the fusion of the IMM between two mitochondria while also serving roles in mitochondrial bioenergetics and cristae architecture ^26^. OPA-1 is just one of several proteins which modulate mitochondrial structure. For example, contrastingly, Dynamin-related protein 1 (DRP-1) is a protein that initiates the fission process through constriction of the mitochondria which divides the mitochondria into two separate organelles ^27^. However, given that OCR increased following insulin treatment, it is possible this is due to the increased mitochondrial area caused by upregulated mitochondrial fusion. To see if OPA-1 may be changed in expression, we performed western blotting. When looking at OPA-1, we noticed a significant continuous increase in protein levels in myoblasts across 2 and 4 hours of insulin stimulation when normalized (**Figure 6A-B**). We further differentiated primary myotubes and carried out these experiments again to see if any differences existed (**Figure 6C-D**). We noticed significant increases in OPA-1 levels after 4 hours of insulin stimulation (**Figure 6C-D**). Together, this suggests that insulin stimulation causes increased expression of OPA-1 in a short time frame which is exacerbated in myotubes compared with myoblasts.

**Figure 6:**
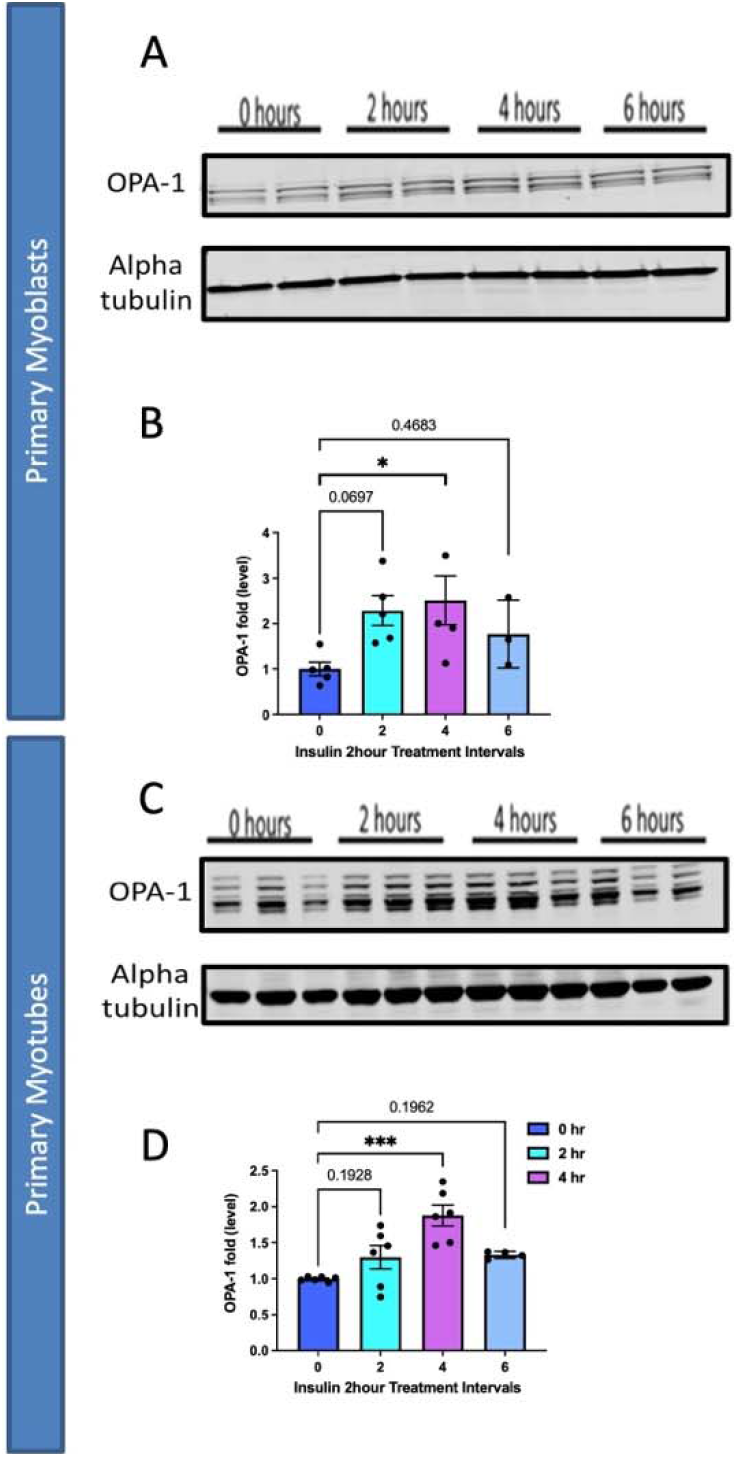
Comparison of mitochondrial fusion proteins following insulin stimulation in primary myoblasts and myotubes. (A) Western blotting for mitochondrial fusion protein OPA-1 following 2 hours, 4 hours, and 6 hours of insulin stimulation. (B) OPA-1 levels normalized to Alpha tubulin following insulin stimulation. (C) This was replicated in primary myotubes, as western blotting for mitochondrial fusion OPA1. (D) OPA-1 levels, normalized to Alpha tubulin, in primary myotubes following insulin treatment. N = 6 per treatment, and * indicates p-value < .05.

Since we saw changes in OPA-1 expression, which is known to trigger fusion, we also validated this technique of myoblasts and myotubes isolation through the quantification of mitochondria and cristae^1^ following insulin treatment. Using transmission electron microscopy, we compared mitochondrial morphology without (**Figure 7A**) and with 2-hour insulin treatment in myoblasts (**Figure 7A’**). When quantified we saw that mitochondria reduce in number (**Figure 7B**), while becoming less spherical and larger in area (**Figure 7C-D**). Together, this suggests an uptick in fusion following insulin treatment in myoblasts. We also considered how cristae morphology may be affected following insulin treatment (**Figure 7E-E’**) and we saw that although the cristae score, a measurement of relative cristae quality, did not change, cristae number and area increased suggesting a greater capacity for oxidative function (**Figure 7F-H**). From there, we similarly sought to see if mitochondria in myotubes had changed following insulin stimulation (**Figure 8A-A’**). Similar to myoblasts, we saw while mitochondria decrease, their size increases (**Figure 8B-D**), again indicating increased mitochondrial fusion. When evaluating the cristae structure, we noticed that while similar to myoblasts cristae score was unchanged, the cristae number had a more significant increase (**Figure 8E-G**). Similar to myoblasts, the cristae area also increased following 2 hours of insulin stimulation (**Figure 8F)**. Together these quantifications show that changes in OCR may be due to OPA-1-mediated changes in mitochondrial and cristae architecture following insulin stimulation.

**Figure 7.**
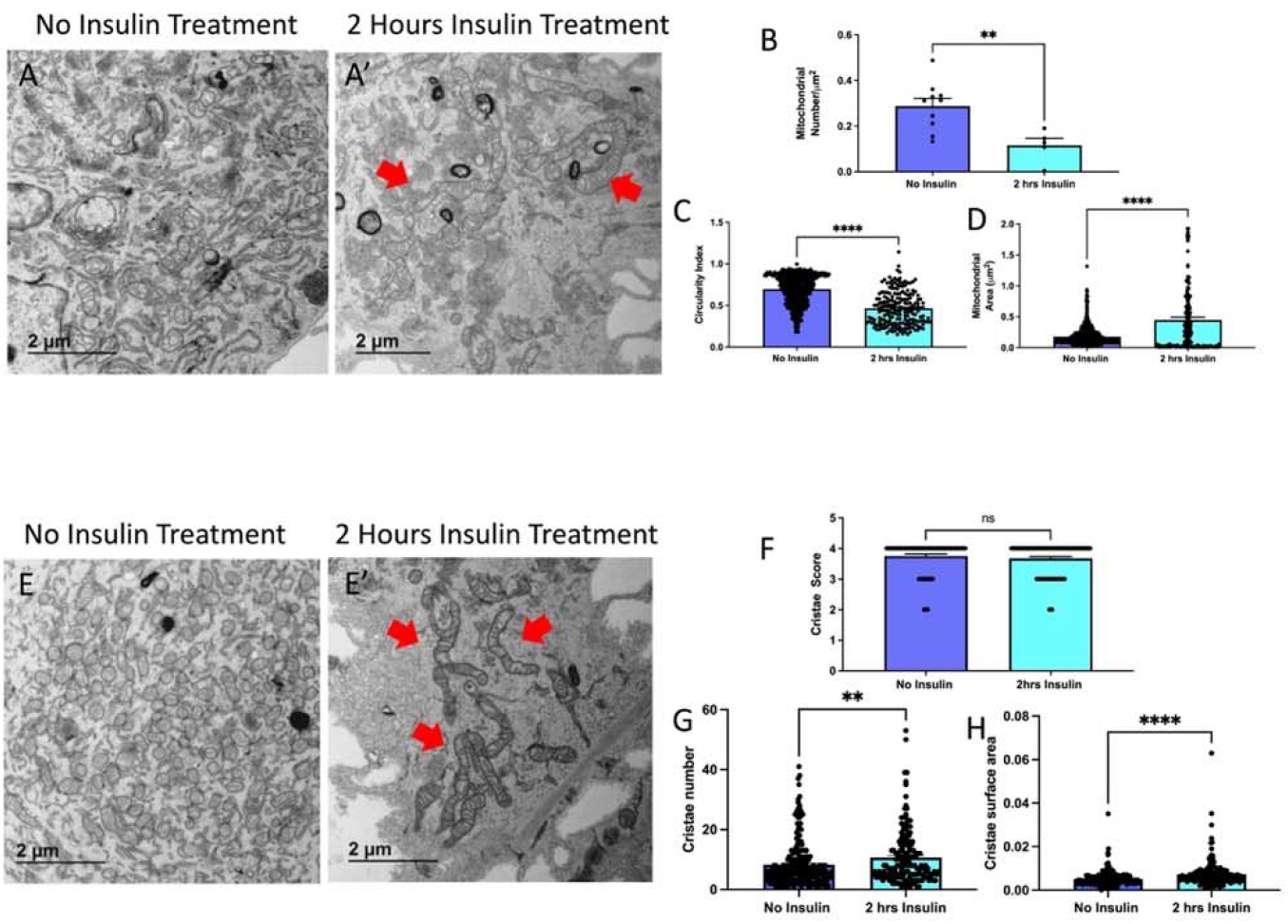
TEM Quantification of Myoblasts. (A) Representative transmission electron micrographs from control and (A’) insulin-treated cells, with red arrows showing fused mitochondria. (B) Quantifications of number of mitochondria, (C) circularity of mitochondria, (D) and area of mitochondria. (E) Representative transmission electron micrographs from control and (E’) insulin-treated cells, with red arrows showing cristae. (F) Quantifications from cristae score, (G) cristae quantity, and (H) cristae area comparing non-insulin and insulin-treated myoblasts. Dots show the number of samples. ** and **** indicates p < 0.01 and p < 0.0001, respectively.

**Figure 8.**
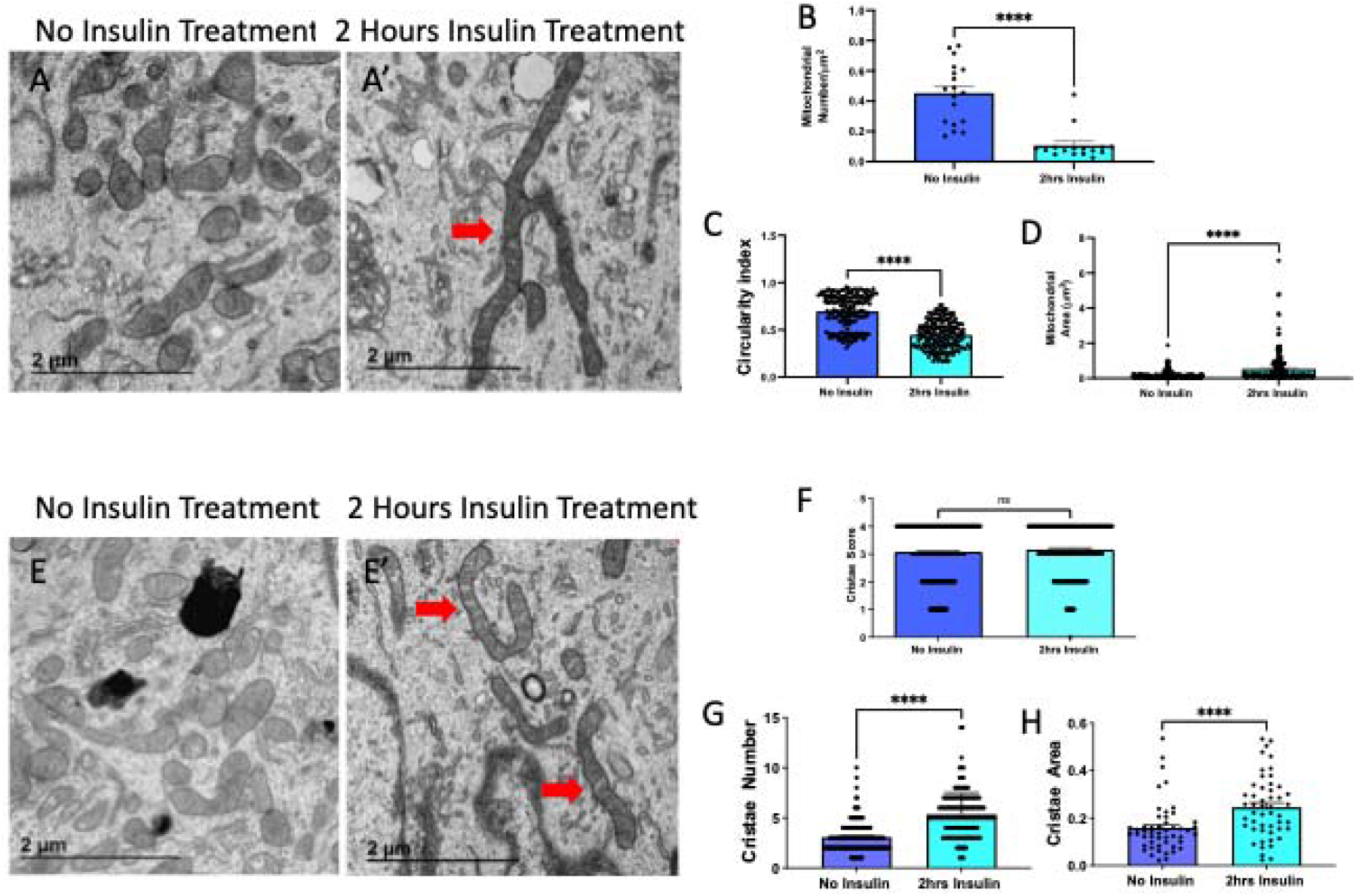
TEM Quantification of Myotubes. (A) Representative transmission electron micrographs from control and (A’) insulin-treated cells, with red arrows showing fused mitochondria. (B) Quantifications of number of mitochondria, (C) circularity of mitochondria, (D) and area of mitochondria. (E) Representative transmission electron micrographs from contro and (E’) insulin-treated cells, with red arrows showing cristae. (F) Quantifications from cristae score, (G) cristae quantity, and (H) cristae area comparing non-insulin and insulin-treated myotubes. Dots show the number of samples. ** and **** indicates p < 0.01 and p < 0.0001, respectively.

Together these data validate this isolation and validation technique allows for the application of experimental models to elucidate cellular processes. This demonstrates the viability of the protocol outlined here for skeletal muscle. After differentiation, quantification can be done for many experimental designs. Here, we performed seahorse analysis per prior methods ^25^ with GraphPad to perform students’ T-tests to measure statistical significance.

Past results have demonstrated that myoblasts can be assayed through staining with Pax7 and MyoD, while multinucleated myotubes are visualized with phase contrast microscopy or after staining for myosin heavy chain ^7^. Past protocols are also available which allow for incredibly dense populations of myoblasts (1.5 Past results have demonstrated that myoblasts × □ 10^7^, 200 times cell expansion) to be obtained through an intelligent culture system with suppression of myotube formation ^12^. Another technique has also allowed for 1 × 10^7^ – 2 × 10^7^ myoblasts to be isolated from murine hindlimbs from a single organism without the need for cell straining or sorting ^28^. Tangentially, techniques allow for induced pluripotent stem cells to be differentiated into myotubes which allow for the study of insulin-resistance ^29^. Our technique is advantageous in offering the ability to obtain either subpopulation, as well as fibroblasts, allowing for a wide range of experiments to perform. Further, the study of specific genes in myoblasts and myotubes can then be affected through verified techniques such as adenovirus, or herpes simplex virus type 1 amplicon vectors, depending on the specific gene ^30^.

## Quantification and Statistical Analysis

For all analyses, GraphPad Prism software package was used (La Jolla, CA, USA), with black bars representing the standard error of meanwhile dots represent individual data points shown. If only two groups were used for comparison, an unpaired t-test was the statistical test, while more than two groups were compared with a one-way ANOVA and Tukey *post hoc* tests for multiple comparisons, or their non-parametric equivalent if applicable. A minimum threshold of *p* < 0.05 indicated a significant difference.

## Limitations

This protocol has been optimized for mice gastrocnemius, quadriceps, and hamstring muscles and may not be applicable to other model organisms or tissue types. Compared with other protocols, ours takes a similar period of time ^28^, but this can still be a slow process that must be carried out across multiple days. While C2C12 myoblasts are ideal for this protocol, increasingly human skeletal myoblasts are important to study, and past protocols indicate that differences in the procedure must be made, such as antisense miR-133a addition, to promote the fast differentiation of human skeletal myoblasts ^31^.

### Trouble Shooting

#### Problem

Ultrastructure or Gross morphology of Myoblasts is Degraded

#### Potential Solution

This may be due to too much damage incurred to myoblasts during preparation. We found that optimizing the process by first digesting tissue with type II collagenase and dispase, followed by grinding the tissue in liquid nitrogen with a mortar with a pestle, and passing it through cell strainers resulted in an improved procedure. However, reducing the time grounded or reducing the amount of digestion can avoid potential damage to the myoblasts if it is occurring.

#### Problem

Contamination with Fibroblasts

#### Potential Solution

It is important to plate first on an uncoated plate. However, if fibroblasts are still observed, pre-plating can be done twice. Antibody-based selection of fibroblasts may cause certain issues but can also be explored as an option to remove fibroblasts. If this remains an issue, other methods have shown that using flowing cytometry can be used to identify and remove fibroblasts ^32^.

#### Problem

Low Cell Yield or Viability

#### Potential Solution

If myoblast or myotube viability is low, it is important to increase the concentration of growth factors and ensure a sterile environment is maintained. Reducing time with accutase can also ensure cells are not treated too harshly.

## Resource Availability

### Lead contact

Further information and requests for resources and reagents should be directed to and will be fulfilled by the lead contact, Antentor Hinton.

### Materials availability

All generated materials, if applicable, are created in methods highlighted in the text above.

### Data and code availability

Full data utilized and requests for data and code availability should be directed to and will be fulfilled by the lead contact, Antentor Hinton.

## Author Contributions

## Acknowledgments

All antibodies were obtained from the Iowa Developmental Studies Hybridoma Bank (DSHB).

## Financial & Competing Interests’ Disclosure

All authors have no competing interests.

This project was funded by the UNCF/Bristol-Myers Squibb E.E. Just Faculty Fund, BWF Career Awards at the Scientific Interface Award, BWF Ad-hoc Award, NIH Small Research Pilot Subaward to 5R25HL106365-12 from the National Institutes of Health PRIDE Program, DK020593, Vanderbilt Diabetes and Research Training Center for DRTC Alzheimer’s Disease Pilot & Feasibility Program. CZI Science Diversity Leadership grant number 2022-253529 from the Chan Zuckerberg Initiative DAF, an advised fund of Silicon Valley Community Foundation (to A.H.J.). NSF EES2112556, NSF EES1817282, and CZI Science Diversity Leadership grant number 2022-253614 from the Chan Zuckerberg Initiative DAF, an advised fund of Silicon Valley Community Foundation (to S.M. D.) and National Institutes of Health grant HD090061 and the Department of Veterans Affairs Office of Research award I01 BX005352 (to J.G.). Additional support was provided by the Vanderbilt Institute for Clinical and Translational Research program supported by the National Center for Research Resources, Grant UL1 RR024975–01, and the National Center for Advancing Translational Sciences, Grant 2 UL1 TR000445–06 and the Cell Imaging Shared Resource.

BioRender was used for the creation of Figures.

## Data Sharing and Open Access

All data is available upon request to the corresponding author.

